# GLUER: integrative analysis of single-cell omics and imaging data by deep neural network

**DOI:** 10.1101/2021.01.25.427845

**Authors:** Tao Peng, Gregory M. Chen, Kai Tan

## Abstract

Single-cell omics assays have become essential tools for identifying and characterizing cell types and states of complex tissues. While each single-modality assay reveals distinctive features about the sequenced cells, true multi-omics assays are still in early stage of development. This notion signifies the importance of computationally integrating single-cell omics data that are conducted on various samples across various modalities. In addition, the advent of multiplexed molecular imaging assays has given rise to a need for computational methods for integrative analysis of single-cell imaging and omics data. Here, we present GLUER (inte<u>G</u>rative ana<u>L</u>ysis of m<u>U</u>lti-omics at single-c<u>E</u>ll <u>R</u>esolution), a flexible tool for integration of single-cell multi-omics data and imaging data. Using multiple true multi-omics data sets as the ground truth, we demonstrate that GLUER achieved significant improvement over existing methods in terms of the accuracy of matching cells across different data modalities resulting in ameliorating downstream analyses such as clustering and trajectory inference. We further demonstrate the broad utility of GLUER for integrating single-cell transcriptomics data with imaging-based spatial proteomics and transcriptomics data. Finally, we extend GLUER to leverage true cell-pair labels when available in true multi-omics data, and show that this approach improves co-embedding and clustering results. With the rapid accumulation of single-cell multi-omics and imaging data, integrated data holds the promise of furthering our understanding of the role of heterogeneity in development and disease.

## INTRODUCTION

A number of single-cell omics assays have been developed for robust profiling of transcriptome, epigenome and 3-dimensional chromosomal organization. Similarly, multiplexed molecular imaging assays have been developed for simultaneous profiling of a large number of proteins ^1,2^ and transcripts ^3–5^ at single-cell resolution. Collectively, these assays provide powerful means to characterize molecular heterogeneity. However, each omics and imaging assay has its own strengths and weaknesses, which results in a partial picture of the biological systems. For example, single-cell omics assays are unable to capture the spatial distribution of measured molecules. On the other hand, imaging assays such as CODEX (co-detection by indexing) ^1^ and MERFISH ^4^ can capture spatial expression patterns of proteins and transcripts within the intricate tissue architecture. However, their coverage is much lower than single-cell omics assays and therefore lack the power to resolve cell types/states ^6^.

Recently, single-cell assays have been developed to jointly measure two or more of molecular modalities in the same cells. For example, sci-CAR ^7^and SNARE-Seq ^8^ allow simultaneous profiling of open chromatin and gene expression. Methyl-HiC ^9^ and single-nucleus methyl-3C ^10^ have been developed to profile chromatin interaction and DNA methylation simultaneously. Although in theory dual-modality measure makes data integration easier, in practice, data integration remains a challenge due to differences in coverage and data modality-specific characteristics.

Computational tools that can flexibly and robustly integrate individual single-cell data sets offer many exciting opportunities for discovery. To date, the most widely used data integration methods are Seurat (v3) ^11^ and LIGER ^12^. Seurat seeks to map data sets into a shared latent space using dimensions of maximum correlations and subsequently maximizes correlated latent factor space which is determined by maximize the correlations. By doing so, it may miss true biological variations across the data sets that may be important. LIGER uses nonnegative matrix factorization and computes the joint clustering using the loading matrices. However, it only considers cell pairs with the smallest distance across the data sets and thus ignore one-to-many and many-to-many pairs, which are biologically meaningful.

Here, we describe GLUER (inteGrative anaLysis of mUlti-omics at single-cEll Resolution by deep neural network). A flexible method for integrating single-cell omics and molecular imaging data. It employs three computational techniques, joint nonnegative matrix factorization, mutual nearest neighbor algorithm, and deep learning neural network. Joint nonnegative matrix factorization of the data sets maintains biological differences across the data sets while allowing identification of common factors shared across the data sets. Mutual nearest neighbor algorithm enables mapping of many-to-many relationships among cells across the data sets. Deep learning neural networks can capture nonlinear relationships between the data sets. In comparison, only linear functions were used in previous methods. Using multiple true multi-omics data sets as the ground truth, we show that GLUER achieved significant improvement in data integration accuracy. We implemented GLUER in Python and also provided a graphical user interface (GUI) for users to explore the integration results.

## RESULTS

### Overview of the GLUER algorithm

GLUER combines joint nonnegative matrix factorization (NMF), mutual nearest neighbor algorithm, and deep neural network to integrate data of different modalities (Figure 1). The purpose of joint NMF is to identify shared components across data sets of different modalities. The result of this step are dimension-reduced matrices (factor loading matrices) for each data modality. The factor loading matrix from one data modality is defined as the reference matrix and the rest as the query matrices. These matrices are used to calculate the cell distance and subsequently determine putative cell pairs between the reference data set and the query data sets. Under the guidance of the putative cell pairs, a deep neural network is used to learn map functions between factor loading matrices of query data sets to the reference factor loading matrix. Using the learnt functions, co-embedded data is then computed by combining the reference factor loading matrix and query factor loading matrices. GLUER is freely available as a Python package at https://github.com/tanlabcode/GLUER.

**Figure 1.**
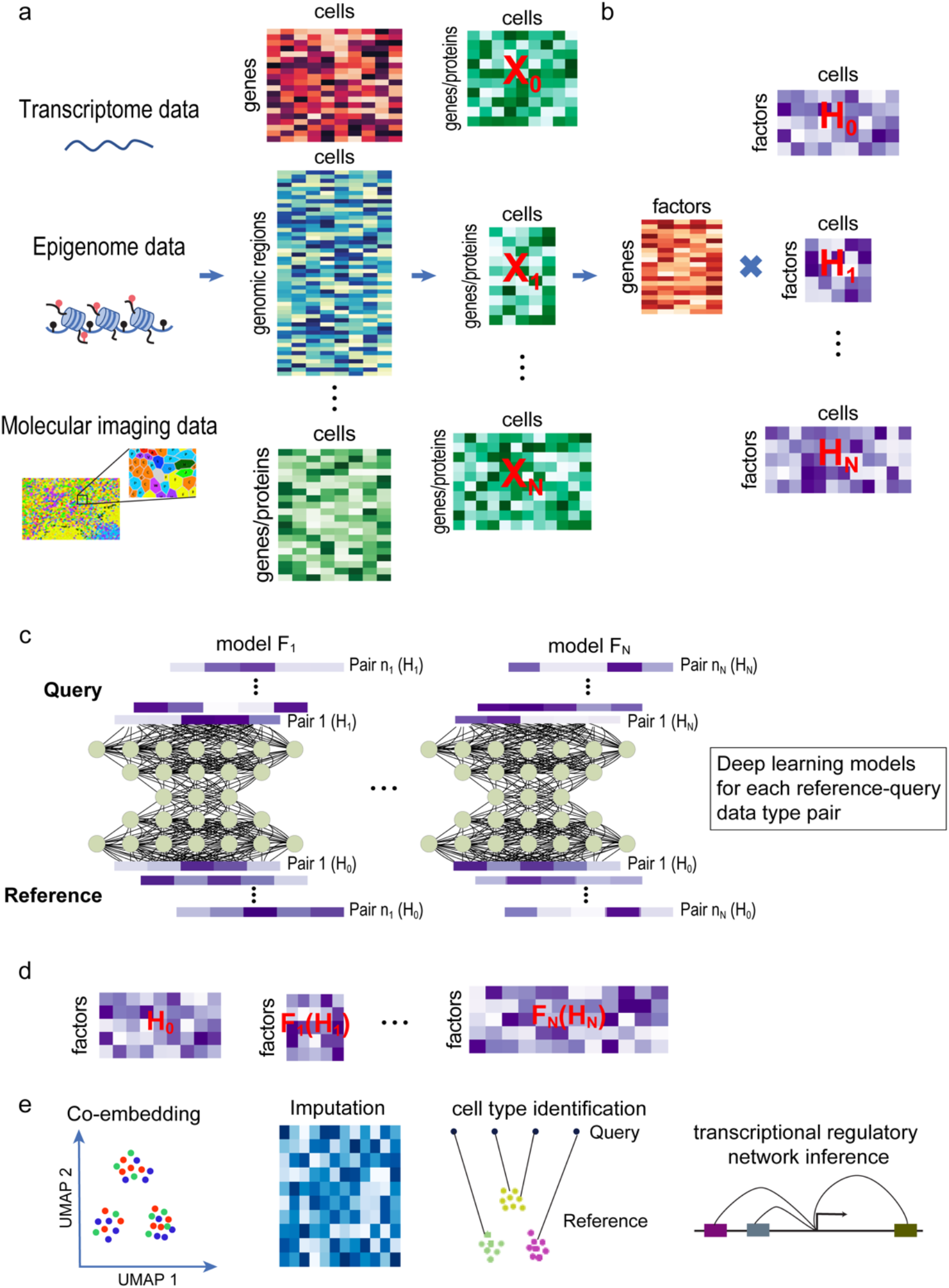
Schematic overview of the GLUER algorithm. (**a**) Representation of single-cell omics and imaging data sets, one of them is the reference modality and the rest are the query modalities. The omics and imaging data sets are processed to generate either gene/protein expression (transcriptomic and imaging data) or gene activity (epigenomic data) data matrices. (**b**) Joint nonnegative matrix factorization to project the normalized data of all modalities into a subspace defined by shared structure across the data sets. Candidate cell pairs between the reference and the query data sets are identified using the mutual nearest neighbor algorithm. (**c**) Deep neural network is employed to determine the nonlinear mapping functions *F_1_, F_2_*,…, *F_N_* from the query data to the reference data based on the cell pairs identified in the previous step. (**d**) The coembedded data matrix is computed using the nonlinear mapping functions. (**e**) Downstream analyses using the co-embedded data, including dimensionality reduction, imputation, cell type identification, and inference of transcriptional regulatory network.

### Performance evaluation using transcriptomics and chromatin accessibility dual-omics data on mixtures of cell lines

We first evaluated the performance of GLUER using true single-cell dual-omics data where the ground truth of cell pairing is known. These data sets were generated using state-of-the-art assays that simultaneously profile mRNA expression and chromatin accessibility, including sci-CAR ^7^, SNARE-Seq ^8^, and scCAT-seq ^13^. sci-CAR and SNARE-Seq process thousands to millions of cells together by using droplet platforms or combinatorial DNA barcoding strategies with high scalability and cost effectiveness. scCAT-seq processes hundreds of single cells in individual wells of microwell plates. Three data sets on mixtures of human cell lines were used (Supplementary Table 1). Data Set 1 was generated using the SNARE-Seq assay ^8^ and a mixture of four human cell lines, BJ, H1, K562, and GM12878. Data Set 2 was generated using the scCAT-seq assay ^13^ and a mixture of 5 human cell lines, including K562, HeLa-S3, HCT116, cells of patient-derived xenograft (PDX) samples of a moderately differentiated squamous cell carcinoma patient (PDX1) and a large-cell lung carcinoma patient (PDX2). Data Set 3 was generated using the sci-CAR assay ^7^ and human lung adenocarcinoma-derived A549 cells after 0,1, and 3 hours of treatment of 100nM dexamethasone (DEX).

We compared three integration methods, Seurat, LIGER, and GLUER using these three data sets, ensuring that each method was blinded to the true cell-pair labels. Thus, this analysis simulated a situation in which single-cell RNA-Seq and ATAC-Seq assays were performed independently on separate batches of cells. We inspected the Uniform Manifold Approximation and Projection (UMAP) plots of the integration results (co-embedded data) and evaluated the results quantitatively using the integration accuracy metric (see Online Methods for details). Since human cell lines were used for generating the ground truth data sets, the true biological identity of the cells is known. Hence, a proper integration should result in retaining the known biological clusters after integrating the data sets.

Figure 2 shows the integration results of Data Set 1 by Seurat, LIGER, and GLUER. A good integration method should generate well-mixed cells identified with different data modalities. Visual inspection of the UMAP plots revealed that the degree of mixing of transcriptome and chromatin accessibility data was the highest in the GLUER result and lowest in the LIGER result (Figure 2a-c). The same trend was also observed in Data Sets 2 and 3 (Supplementary Figures 1a-c, 2a-c). By design, the cell line mixture data provides the ground truth for the number of cell clusters we should expect in the integrated data. For Data Set 1, LIGER result produced 10 cell clusters, which is not consistent with the fact that only four cell lines were used for generating the SNARE-Seq data set ^5^ (Figure 2b). Similar result was also observed with Data Set 2 where LIGER result yielded more than 5 cell clusters which was the number of cell lines used to generated the scCAT-Seq data set ^3^ (Supplementary Figure 1b). Seurat result yielded the correct number of clusters (four) on Data Set 1 although separation between two of the clusters was less clear (Figure 2d). Only GLUER result yielded four cell clusters that were well separated and represented the four cell lines in the mixture sample (Figure 2f). For Data Set 3, only GLUER result can separate DEX untreated and treated A549 cells (Supplementary Figure 2d-f). Finally, the expression patterns of known marker genes for the cell lines confirm their identities (Figure 2i, Supplementary Figure 1h).

**Figure 2.**
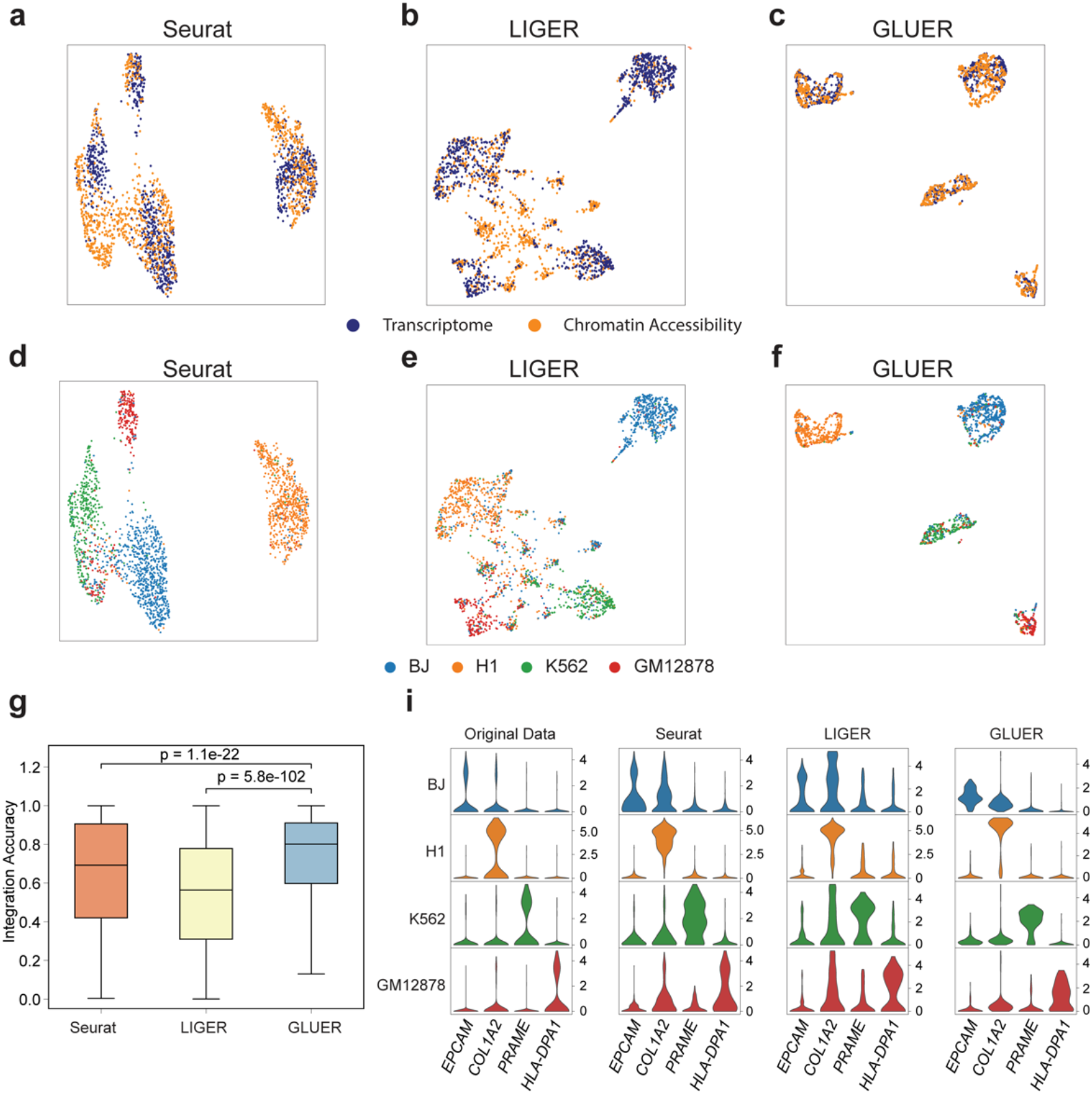
Performance evaluation using data on a mixture of 1047 human BJ, H1, K562 and GM12878 cells profiled using SNARE-Seq. (**a-c**) UMAP plots of integrated datasets, color-coded by data modality. (**d-f**) UMAP plots of integrated datasets, color-coded by cell line. (**g**) Integration accuracy of the three methods. P-values were computed using paired Student’s *t*-test with Bonferroni correction. (**i**) Distributions of gene expressing levels in the original scRNA-Seq data and data computed by the integration methods. Y-axis, normalized gene expression level. *EPCAM, COL1A2, PRAME*, and *HLA-DPA1* are the marker genes for BJ, H1, K562, and GM12878 cell lines, respectively.

To quantitatively evaluate the performance of the methods, we devised a metric called integration accuracy that is the percentile of the true cell pairs among the predicted neighbors of a cell. It is calculated based on the dimension-reduced matrices of the co-embedded data (see Online Methods for details). This metric ranges from 0 to 1 and is high when the true cell pairs across the data sets share a high percentage of the same neighbors. Our analysis indicates that GLUER had significantly higher integration accuracy compared to Seurat and LIGER for all three data sets (p < 0.05, paired Student’s *t*-test; Figure 2g, Supplementary Figures 1g, 2g).

### Performance evaluation using transcriptomics and chromatin accessibility dual-omics data on primary tissues

To further evaluate the performance of GLUER on data of complex tissues, we used three true dual-omics data sets of primary tissue samples generated using SNARE-Seq and sci-CAR assays (Supplementary Table 1). Data Sets 1 and 2 ^8^ were generated using SNARE-Seq and neonatal mouse cerebral cortex and adult mouse cerebral cortex samples, respectively. Data Set 3 ^7^ was generated using sci-CAR and mouse kidney sample.

Figure 3 shows integration results of Data Set 1 by Seurat, LIGER, and GLUER. Visual inspection of the UMAP plots revealed that the degree of mixing of transcriptome and chromatin accessibility data was the highest in the GLUER result and lowest in the Seurat result (Figure 3a-c). The same trend was also observed in Data Sets 2 and 3 (Supplementary Figures 3a-c, 4a-c). Clustering analysis using the Leiden method ^14^ revealed more cell clusters in GLUER result versus results of the other two methods, which is consistent with fact of large numbers of cell types in these tissues (Figure 3e, Supplementary Figures 3e and 4e). For instance, in Data Set 1, integrated data by GLUER revealed 14 cell clusters, representing excitatory neurons in Layer 2/3 (EL-2/3-Rasgrf2), Layer 3/4 (Ex-L3/4-Rorb), Layer 4/5 (Ex-L4/5-Il1rapl2 and Ex-L4/5-Epha4), Layer 5 (Ex-L5-Klhl29 and Ex-L5-Parm1), Layer 5/6 (Ex-L5/6-Tshz2 and Ex-L5/6-Sulf1), and Layer 6 (Ex-L6-Foxp2), inhibitory neurons (In-Npy, In-Sst, and In-Pvalb), claustrum, and astrocytes/oligodendrocytes (Ast/Oli) (Figure 3e). Interestingly, the different types of excitatory neurons were closer to each other in the UMAP, reminiscent of their adjacent physical locations in the mouse cerebral cortex. The identification of each cell cluster was determined based on gene expression and gene activity profiles of known cell-type markers, which was calculated using the co-embedded transcriptome and chromatin accessibility data. The marker gene expression and activity profiles clearly demonstrate cell type specificity (Figure 3f and 3g). More cell clusters were also identified in Data Sets 2 (adult mouse cerebral cortex, Supplementary Figure 3e-g) and 3 (mouse kidney, Supplementary Figure 4e-g) integrated by GLUER.

**Figure 3.**
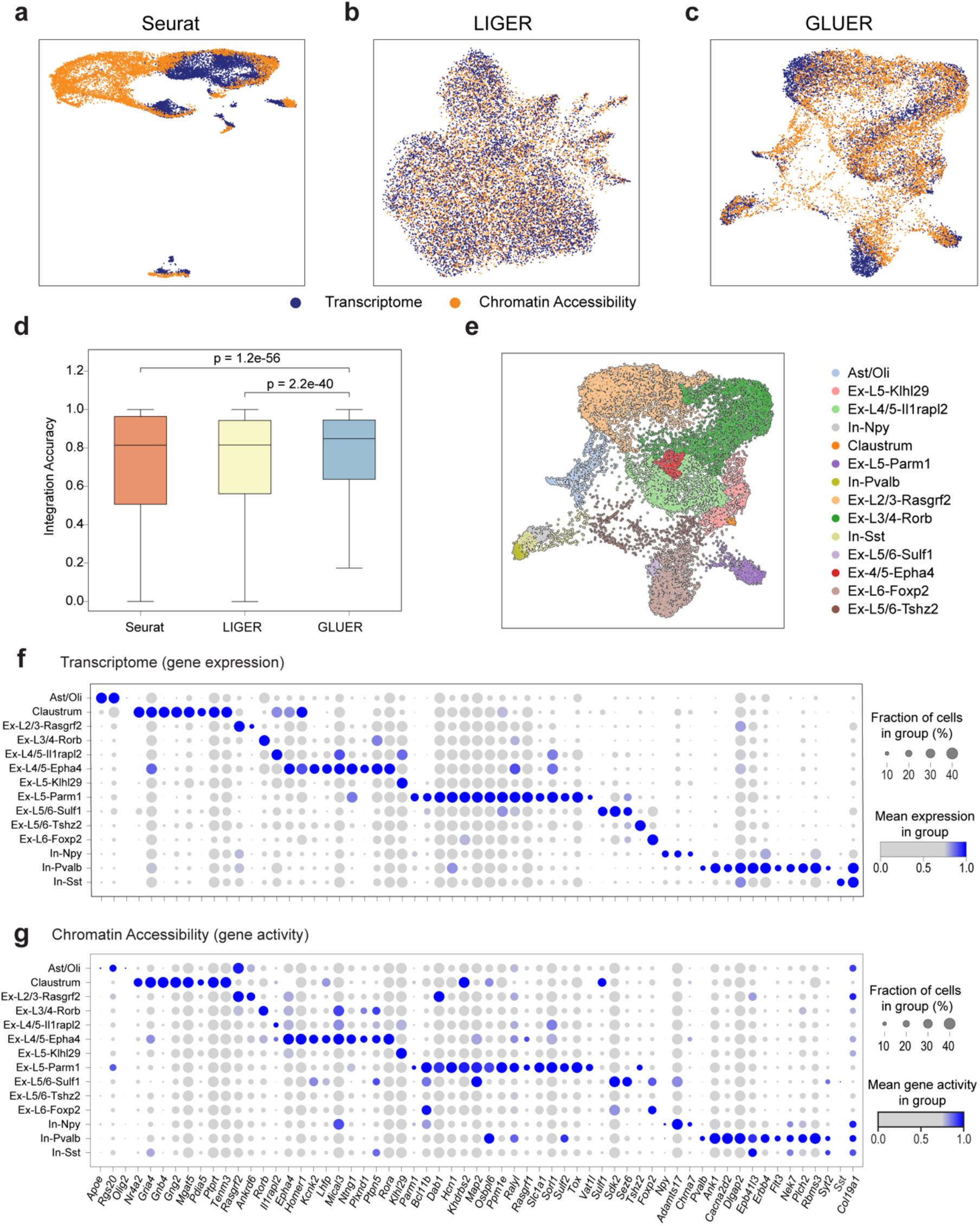
Performance evaluation using data on adult mouse cerebral cortex. 7,892 cells were profiled using SNARE-Seq. (**a-c**) UMAP plots of integrated datasets, color coded by data modality. (**d**) Integration accuracy of the three methods. P-values were computed using paired Student’s *t*-test with Bonferroni correction. (**e**) UMAP plot of integrated data by GLUER with cell type annotation. Ex-L2/3, excitatory neurons in Layer 2/3, the same for the rest of excitatory neuron abbreviation; In, inhibitory neurons; Ast, astrocytes; Oli: Oligodendrocytes. (**f**) Marker gene expression profiles of the cell types identified in panel e. (**g**) Marker gene activity profiles of the cell types identified in panel e. Y-axis, cell types, X-axis, marker genes. Color of the dots represents normalized gene expression (for RNA-Seq) or gene activity (for ATAC-Seq). Size of the dots represents the percentage of cells with nonzero expression or activity of the gene.

Using the integration accuracy metric, we quantified the performance and found that GLUER has the highest accuracy in all three data sets (p < 0.05, paired Student’s *t*-test; Figure 3d and Supplementary Figures 3d and 4d).

In summary, using both cell line mixture and primary tissue data where the ground truth of cell-pairs is known, we demonstrate that GLUER achieved significant improvement in integrating single-cell transcriptomic and chromatin accessibility data.

### Use case 1: Integrating spatial proteomics data with scRNA-Seq data

Co-detection by indexing (CODEX), a multiplexed cytometric imaging method, can simultaneously profile the expression levels of several dozens of proteins in a tissue section with single-cell resolution ^6^. However, the ability of the method to resolve cell types/states is limited due to the small number of features (proteins) in a typical data set. On the other hand, scRNA-Seq assays can profile the expression levels of thousands of genes in each cell, which enables much finer classification of cell types and states. To demonstrate the utility of GLUER for integrating spatial proteomics data and single-cell omics data, we applied it to a matched data set of murine spleen cells that consists of 7,097 cells profiled by scRNA-Seq ^15^ and 9,186 cells profiled by CODEX ^6^ (Supplementary Table 1). A 30-antibody panel was used in the CODEX experiment to identify splenic-resident cell types.

Inspecting the UMAP plots of the original scRNA-Seq data and CODEX data, multiple cell clusters are readily evident in the scRNA-Seq data whereas cluster structure is not clear in the CODEX data (Figure 4a). After data integration and clustering of the co-embedded data using the Leiden method, cluster structures in the UMAP plots are more evident in the Seurat and GLUER results than that of the LIGER result (Figure 4a). 17, 20, and 18 clusters were identified based on the integrated results by Seurat, LIGER and GLUER respectively (Figure 4b). Using known cell-type marker genes, we annotated the clusters based on GLUER integration result (Figure 4c and Supplementary Figure 5). GLUER identified NK cells, red pulp macrophages, neutrophils, monocytes, pDCs, plasma cells, erythrocytes/erythroblasts, B cells, and T cells (Figure 4c). Integrated result by GLUER can effectively separate distinct cell types, for instance NK cells and red pulp macrophages. The latter are distinct from monocytes and monocyte-derived macrophages. Natural cytotoxicity triggering receptor (*Ncr1*) ^16^ and vascular cell adhesion molecule 1 (*Vcam1*)^17^ are specific markers for these two cell types, respectively. In the original data, *Vcam1* transcript expression was restricted to a cluster of cells in the scRNA-Seq UMAP but VCAM1 protein expression was more scattered in the CODEX UMAP. In the integrated data, both transcript and protein expression were mostly restricted in one cluster in the GLUER result (cluster 12) compared to more scattered expression in Seurat (centered around clusters 6 and 12) and LIGER (centered around clusters 1 and 18) results (Figure 4d and 4f). Similar to *Vcam1/VCAM1*, *Ncr1/NCR1* expression was restricted to a cluster of cells in the scRNA-Seq UMAP but more scattered in the CODEX UMAP. In the integrated data, both transcript and protein expression were mostly restricted in one cluster in the GLUER result (cluster 16) compared to Seurat (centered around clusters 6 and 12) and LIGER (centered around clusters 1 and 17) results (Figure 4e and 4g). In summary, we show that by integrating with scRNA-Seq data, cell types that are hard to distinguish in the original CODEX data can be readily identified. Using the co-embedded data, spatial distribution of additional genes that are not covered in the CODEX panel can then be studied.

**Figure 4.**
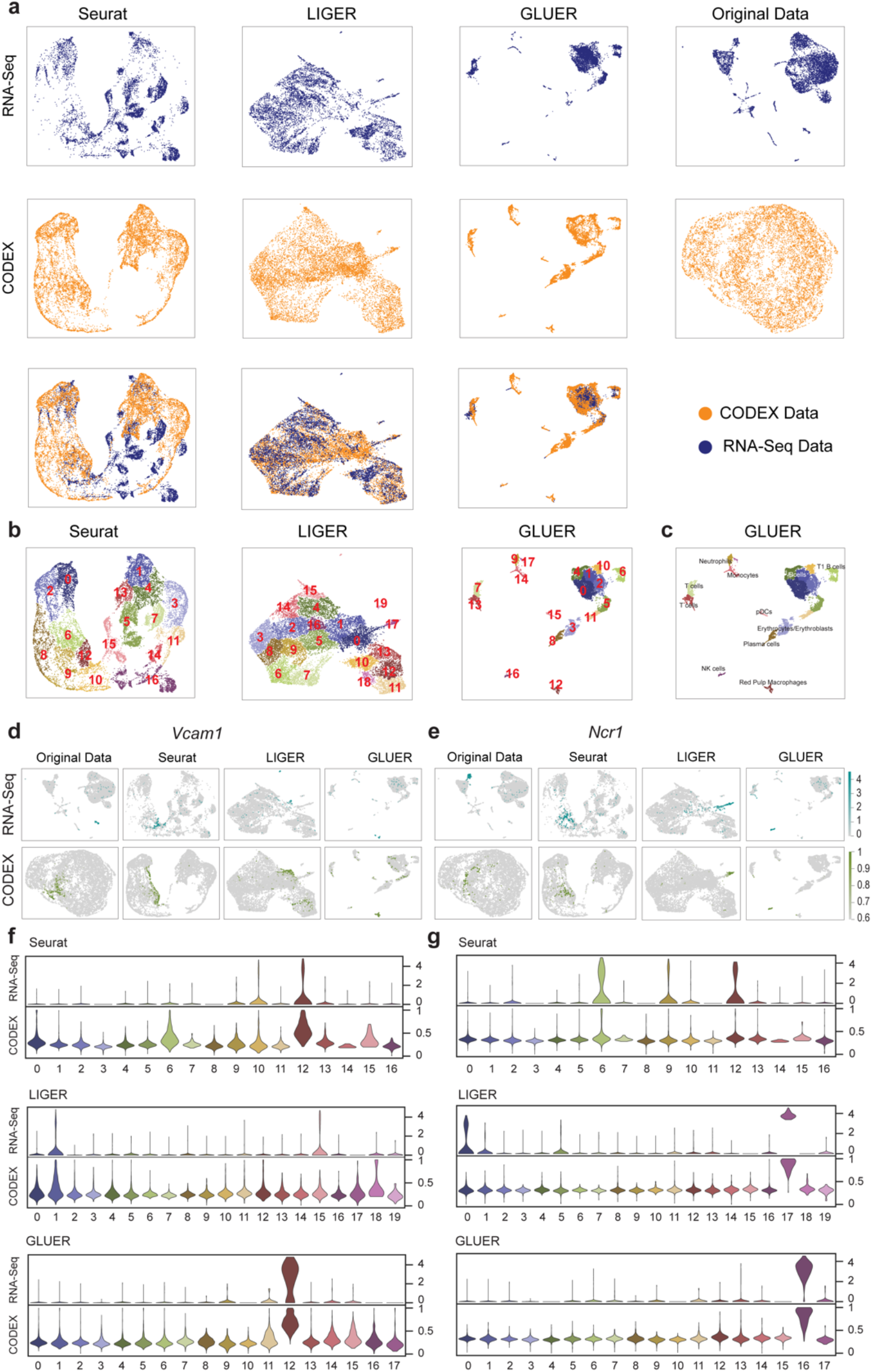
Integration of scRNA-Seq and single-cell spatial proteomics data. The dataset consists of scRNA-Seq and CODEX data on mouse spleen. (**a**) UMAP plots of the integrated data and original data. First row, UMAPs based on the scRNA-Seq part of the co-embedded data; Second row, UMAPs based on the CODEX part of the co-embedded data; Third row, UMAPs based co-embedded scRNA-Seq and CODEX data. (**b**) Cell clusters identified using integrated data by Seurat, LIGER, and GLUER. Clustering was performed using the Leiden method. (**c**) Cell type annotation based on data integrated using GLUER. (**d-e**) UMAP of marker gene/protein expression for red pulp macrophages and NK cells. *Vcam1*, marker gene for red pulp macrophages; *NKp46*, marker gene for NK cells. (**f-g**) Violin plots of *Vcam1* and *NKp46* expression levels across cell clusters. Y-axis, normalized expression levels of genes/proteins.

### Use case 2: Integrating spatial transcriptomics data and scRNA-Seq data

Highly multiplexed single-molecule fluorescence *in situ* hybridization (smFISH) assays such as sequential fluorescence *in situ* hybridization (SeqFISH+) ^3^ and multiplexed error-robust fluorescence *in situ* hybridization (MERFISH) ^4^ enable spatial profiling of hundreds to thousands of transcripts at single-cell resolution. Here we demonstrate the utility of GLUER for integrating scRNA-Seq data with SeqFISH+ data. We used matched data sets on mouse visual cortex and olfactory bulb ^3, 18^, respectively (Supplementary Table 1). The scRNA-Seq data set consists of 1344 and 17709 cells for the two tissue types and the SeqFISH+ data set consists of 523 and 2050 cells for the two tissue types. Similar to CODEX data, due to the smaller number of features in the SeqFISH+ data, UMAP plot of the original data does not reveal clear clusters (Figure 5c,d and 5g,h). After applying GLUER, UMAP plots show that cells identified by both assays are well mixed. Clustering of the co-embedded data revealed 5 major clusters representing oligodendrocytes, endothelial cells, astrocytes, GABAergic neurons, and glutamatergic neurons (Figure 5a and 5b). In general, the same cell types labeled by both data modalities clustered together. For instance, oligodendrocytes, endothelial cells, and astrocytes from SeqFISH+ data and scRNA-Seq data clustered together. L5a, L5b, L2/3, L6a, L4_Ctxn3/Scnn1a neurons from SeqFISH+ data and glutamatergic neurons from scRNA-Seq data clustered together. Pvalb, L6b, Sst, Lgtp/Smad3 neurons from SeqFISH+ data and GABAergic neurons from scRNA-Seq clustered together. Using a second data set on mouse olfactory bulb ^3, 19^, we show that GLUER yielded similar result (Figure 5e-h). Again, UMAP plot of the original SeqFISH+ data did not reveal clear clusters in the SeqFISH+ data (Figure 5h). After data integration, many cell types labeled by both modalities were clustered together, such as astrocytes, microglia cells and macrophages, and endothelial cells. In summary, our result demonstrates that GLUER is able to accurately integrate multiplexed RNA FISH data with scRNA-Seq data.

**Figure 5.**
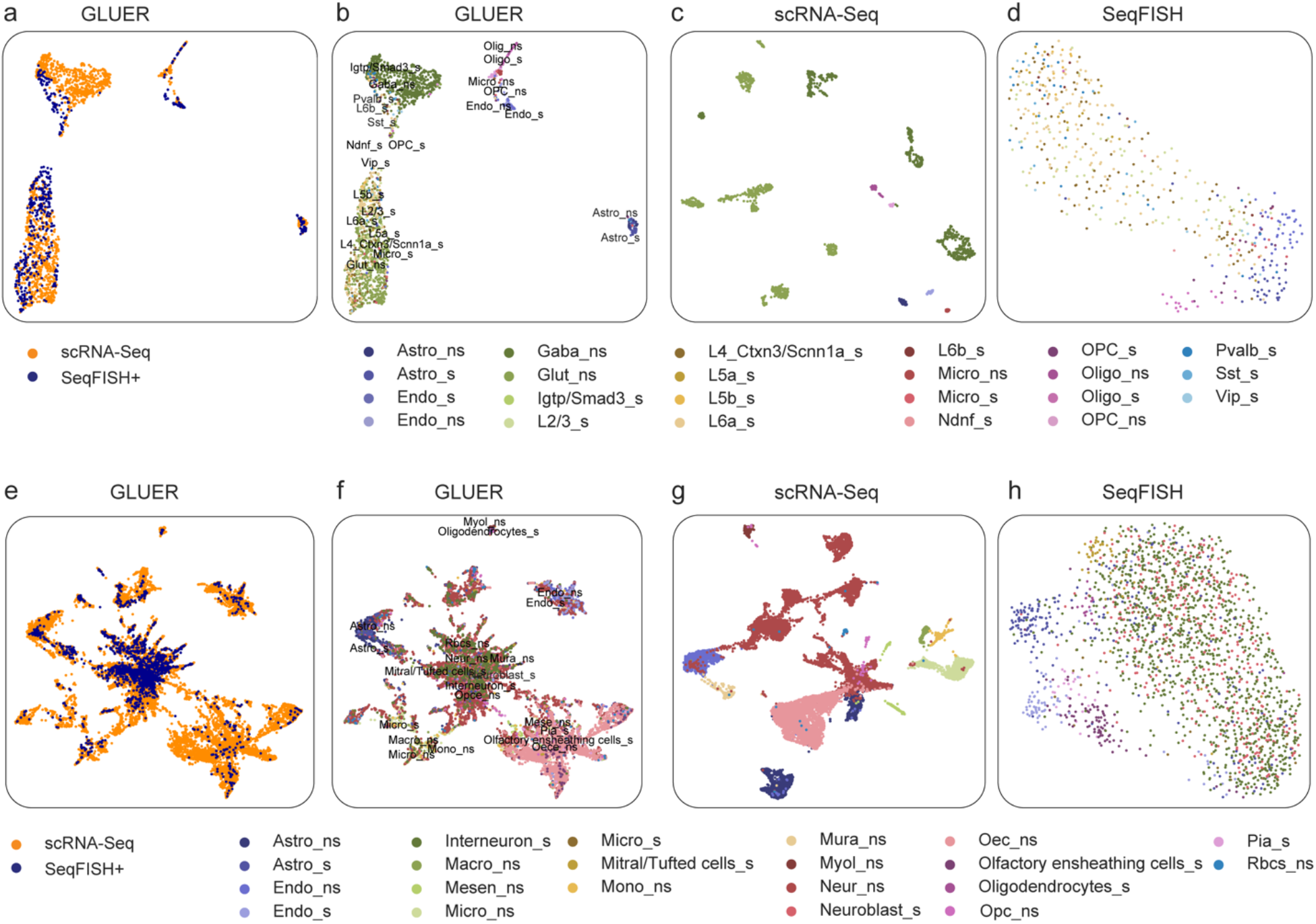
Integration of scRNA-Seq and single-cell spatial transcriptomics data. (**a-d**) UMAP plots of mouse visual cortex data. (**a**) Integrated data annotated by data modalities, SeqFISH+ and scRNA-Seq. (**b**) Integrated data annotated by cell types. Cell type annotation is from original publications. Cell types from scRNA-Seq and SeqFISH+ data are labeled with “ns” and “s”, respectively. Astro, astrocytes; Endo, endothelial cells; Gaba, GABAergic neurons; Lgtp/Smad3, Interneurons highly expressing Lgtp/Smad3; Pvalb, Interneuron highly expressing Pvalb; Sst, Interneurons highly expressing Sst; Vip, Interneuron highly expressing Vip; Ndnf, Interneurons highly expressing Ndnf; Glut, glutamatergic neurons; L2/3, excitatory neurons in Layer 2/3; L4_Ctxn3/Scnn1a, excitatory neurons in Layer 4 highly expressing Ctxn3/Scnn1a; L5a, excitatory neurons in Layer 5a; L5b, excitatory neurons in Layer 5b; L6a, excitatory neurons in Layer 6a; L6b, excitatory neurons in Layer 6b; OPC, oligodendrocyte precursor cells; Oligo, oligodendrocytes; Micro, microglia cells; (**c**) scRNA-Seq data alone (**d**) SeqFISH+ data alone. (**e-h**) UMAP plots of mouse olfactory bulb data. (e) Integrated data annotated by data modalities. (f) Integrated data annotated by cell types. (g) scRNA-Seq data alone. (h) SeqFISH+ data alone. Astro, astrocytes; Endo, endothelial cells; Micro, microglia cells; Neur, neurons; Myol, myelinating-oligodendrocytes; Mesen, mesenchymal cells; Mura, mural cells; Mono, monocytes; Macro, macrophages; Rbcs, red blood cells; Opc, oligodendrocyte progenitor cells; Oec, olfactory ensheathing cells; Pia, pial cells;

### Use case 3: Improving integration of true transcriptomics and epigenomics dual-omics data

True dual/multi-omics protocols have started to emerge recently. Although the different modalities are measured in the same cells, integration of such data remains a challenge due to differences in coverage and data modality-specific characteristics. For instance, transcriptomic data in dual scRNA-Seq and scATAC-Seq data typically reveals more granular structure in the data, as demonstrated using a SNARE-Seq data set on adult mouse cerebral cortex (Figure 6a, 6b) (Supplementary Table 1). Here, we illustrate the ability of GLUER to improve integration of true dual/multi-omics data. One of the key steps of GLUER is inference of cell pairs among different data modalities using mutual nearest neighbor algorithm (Figure 1b). With true dual/multi-omics data, instead of inferring cell pairs using the mutual nearest neighbor algorithm, GLUER can take advantage of the true cell-pair information to perform joint dimensionality reduction and co-embedding. After integration of the mouse cerebral cortex SNARE-Seq data set, cells labeled by the two data modalities were well mixed (Figure 6c). Clustering analysis of the integrated data uncovered 15 clusters (Figure 6e) that can be annotated to known cerebral cortex cell types based on known marker genes. Importantly, based on the co-embedded data, expression profiles of marker genes across cell types are consistent with gene activity profiles across cell types (Figure 6f and 6g), confirming the high degree of mixing of cells identified by these two data modalities in the UMAP plot.

Compared to Seurat and LIGER, the integration accuracy of GLUER is significantly higher (p < 0.05, paired Student’s *t*-test; Figure 6d).

**Figure 6.**
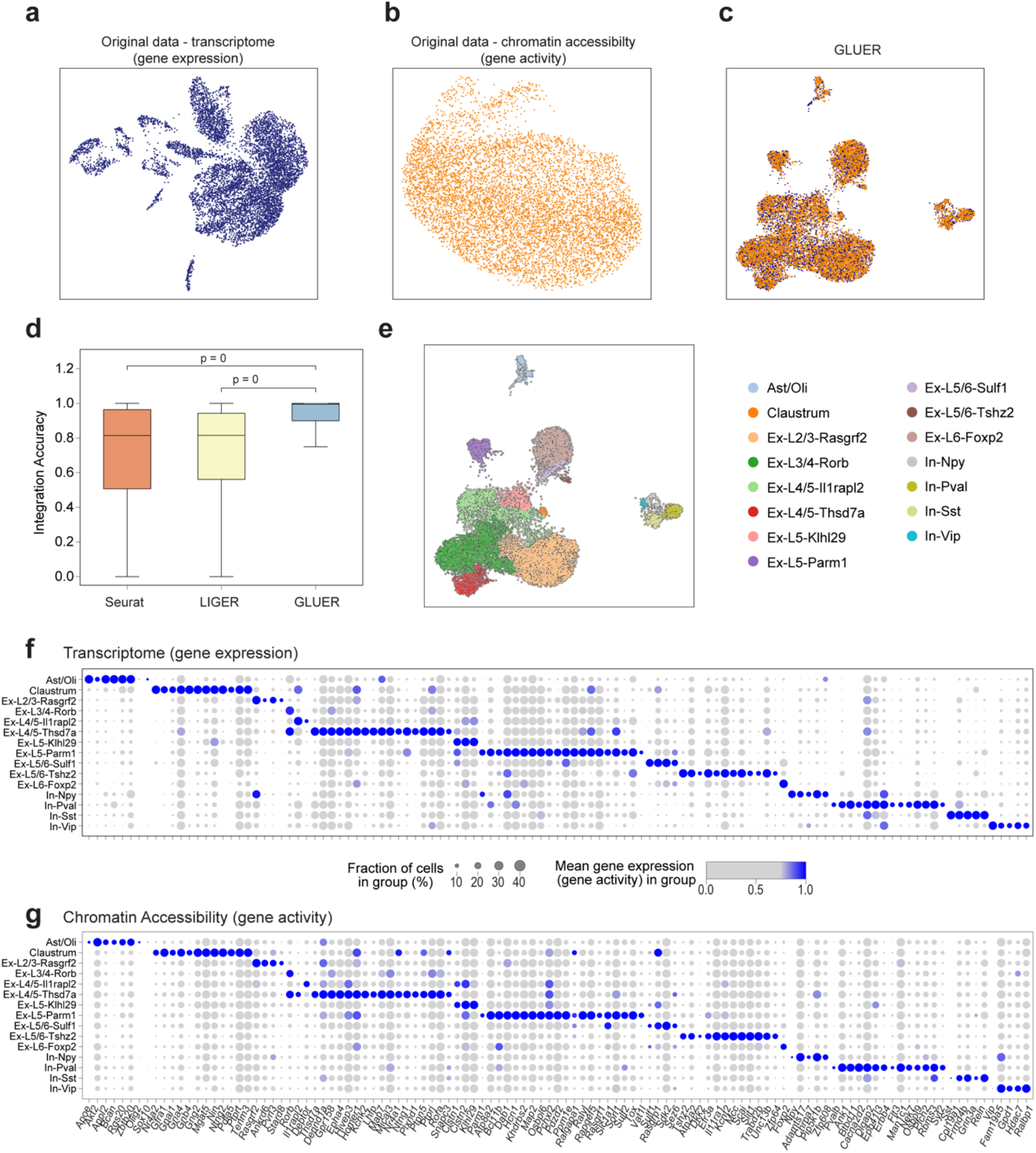
GLUER improves integration of true single-cell dual-omics data. The data set consists of 7,892 cells from adult mouse cerebral cortex profiled using SNARE-Seq. (**ac**) UMAP plots of original data and integrated data by GLUER. (**d**) Integration accuracy of the three methods. P-values were computed using paired Student’s *t*-test with Bonferroni correction. (**e**) UMAP plot of integrated data by GLUER with cell type annotation. Ast, astrocytes; Oli, oligodendrocytes; Ex-L2/3-Rasgrf2, excitatory neuron highly expressing *Rasgrf2*; In-Npy, inhibitory neuron highly expressing *Npy*. The same nomenclature applies to other excitatory and inhibitory neuron subtypes. (**f**) Marker gene expression profiles of the cell types identified in panel e. (**g**) Marker gene activity profiles of the cell types identified in panel e. Y-axis, cell types, X-axis, marker genes.

## DISCUSSION

An accurate definition of cell type/state and underlying gene regulatory mechanisms requires integration of multiple measurement modalities at the single cell resolution. GLUER was designed as a general framework for single-cell data integration. Integration can be performed for multiple single-modality omics data sets, single-modality omics data and multiplexed molecular imaging data, and different measurement modalities of true multi-omics data. Each of the three major steps of GLUER addresses a specific challenge in data integration. The joint nonnegative matrix factorization addresses the high dimensionality of single-cell omics data. The mutual nearest neighbor algorithm can capture many-versus-many relationships among cells and thus further reduce the search space for cells of matched data modalities. The deep neural network can capture non-linear functional relationships between different data modalities. The combination of these three critical steps enables robust integration of high-dimensional and multi-modality data. Using diverse single-cell data sets generated with different protocols and tissue types, we demonstrated that GLUER achieved significantly higher integration accuracy over existing methods.

The GLUER software offers additional technical benefits. First, the software is high scalable due to its implementation of parallel computing using CPUs and GPUs. It takes <5 minutes to integrate ~10,000 cells with multi-omics data. Second, GLUER is built on the AnnData object which supports preprocessing and plotting functions in SCANPY ^20^. Third, the graphical user interface (GUI) allows interactive exploration of the integration results.

Here, in order to conduct joint NMF of transcriptomic/proteomic data and epigenomic data, we summarized the epigenomic data (chromatin accessibility data in this case) as a gene activity matrix. For each gene, its activity score is the sum of reads in its promoter region and gene body regions. Novel approaches for computing gene activity matrices that take into account additional information located in distal regulatory regions in the genome may help to improve integration accuracy. Compared to chromatin accessibility, the relationship between DNA methylation and gene expression is more complicated. More sophisticated functions other than summation are needed to accurately and comprehensively capture the relationship of DNA methylation and gene expression to construct a gene activity matrix.

In summary, GLUER is a much-needed tool for integrating single-cell data across measurement modalities and/or samples. With the rapid accumulation of single-cell multi-omics and imaging data, integrated data holds the promise of furthering our understanding of the role of heterogeneity in development and disease.

## ONLINE METHODS

### Overview of GLUER

GLUER integrates single-cell omics and molecular imaging data using three major computational techniques: joint nonnegative matrix factorization, mutual nearest neighbors, and convolutional deep neural network. Specifically, it consists of the following steps (Figure 1): (a) normalize data for each data modality; (b) perform joint nonnegative matrix factorization of the different data sets and identify cell pairs using the mutual nearest neighbor algorithm; (c) learn a nonlinear mapping function between the factor loadings of a pair of data modalities using convolutional deep neural network; (d) generate a co-embedding matrix and jointly visualize the cells using UMAP or t-SNE with the option of performing imputation on the co-embedded data.

### Normalization of data of different modalities

For transcriptomic data, we used a cell-by-gene matrix of unique molecular identifier (UMI) counts as the input for each algorithm. For chromatin accessibility data, we generated a gene activity matrix by aggregating UMI counts in the 2.5 kb upstream region of transcription start site and the gene body of each gene. For other types of omics data, we could use the commonly used approaches to generate the gene activity matrix. We employed the same standard pre-processing procedures to generate gene expression matrices and gene activity matrices from all data sets used in this paper. We applied a log-normalization of the gene expression matrices or gene activity matrices of all data sets using a size factor of 10,000 molecules for each cell. We removed low-quality cells before normalization. For CODEX data, single-cell protein expression levels were calculated using the average intensity of the pixels constituting each single cell. We first removed cells smaller than 1,000 or larger than 25,000 voxels (the default settings of STvEA ^15^). Then we removed noise in the CODEX data by fitting a Gaussian mixture model to the expression levels of each protein ^9^ followed by normalizing the data by the total levels of protein markers in each cell. The output of the normalization step of GLUER are normalized gene expression, gene activity, and protein expression matrices denoted as *X_0_,X_1_*,…, *X_N_*, where *X*_0_ is the reference data set. For each *x_i_* (*i* = 0,…,*N*), rows are genes/proteins and columns are individual cells. All *X_i_* (*i* = 0,1,…,*N*) have the same features (gene/protein) and we use *g* to denote the number of features and *m*_0_,*m*_1_,…,*m_N_* to denote the number of the cells in *X_i_* (*i* = 0,1,…,*N*) respectively.

### Joint nonnegative matrix factorization of multiple datasets

We use the following model to compute the joint non-negative matrix factorization

(NMF) of the multiple data sets. Given normalized data matrices *X_i_,i* = 0,…,*N*, we can compute the factor loadings matrices *H_0_*,*H_1_*,…,*H_N_* for each data set by minimizing the following cost function:

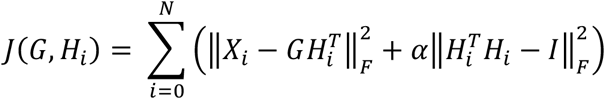

where 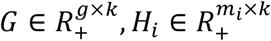 (*k* is the number of factors in the joint NMF) and *I* is the identity matrix. The first term in the sum represents approximation error and the second term the orthogonality of column vectors in *H_i_*, where the trade-off is controlled by the hyperparameter *α*. The optimization problem is non-convex, which means no global optimal solution for the problem. However, it can be solved by the following iterative scheme to reach its local minimum. The algorithm starts by initializing the values in *G* and *H_i_* randomly and uniformly distributed on [*0,1*], and updating them with the following rules until convergence.

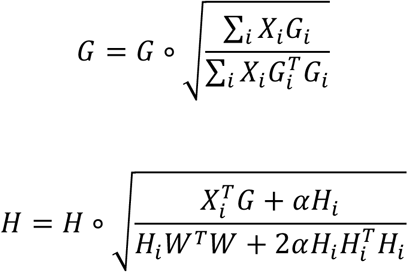

where ° represents the element-wise (Hadamard) product ^21^. The mathematical proof of the iterative procedure and its convergence to the stationary points have been shown in previous literature ^21, 22^.

### Identification of candidate cell pairs using mutual nearest neighbor algorithm

We identify candidate cell pairs between the reference data (*H_0_*) and query data (*H_1_*, *H_N_*) using the mutual nearest neighbor (MNN) algorithm where cell distance is determined by the cosine distance between *H_0_* and *H_1_*,….,*H_N_*. Given that scRNA-Seq data has highest coverage among the data modalities and the central role of transcription linking epigenomic and protein information, by default, scRNA-Seq is chosen as the reference data modality. The MNN algorithm was first introduced for removing batch effects and it is also implemented by Seurat ^11, 23^. Some cells may appear in a large number of cell pairs. These dominant cells are more likely to appear in the integrated data. Furthermore, we found that the number of dominant cells is highly context- and sample-dependent. Here we employ the following steps to reduce the bias due to the dominant cells while identifying cell pairs. We use introduce two tunable parameters for identifying dominant cells. First, we check if cells of a candidate cell pair are also mutual nearest neighbors in the high dimensional space whose distance is calculated directly from the normalized data. The distance calculation can be tuned by using the joint_rank parameter, which is the number of shared factors in the joint nonnegative matrix factorization step. Second, for each cell, we keep its top K nearest neighbors in the pairs. Hence, the algorithm is seeking K mutual nearest neighbors.

### Finding nonlinear mappings among the factor loadings of different datasets using a deep neural network

In order to project data from different biological assays onto a common feature space, we investigated functions *F_i_, i* ∈ {*1,….N*} to map the factor loadings *H_1_*,….,*H_N_* onto the feature space of *H_0_*. Previous methods such as Seurat uses a linear function. In contrast, we use a convolutional deep neural network to capture nonlinear relationships among the factor loading matrices of different modalities (*H_i_* and *H_0_*) based on the identified cell pairs *P_i_*. *P_i_* is a matrix consisting of two columns, the first column is the cell index in the reference data set and the second column is the cell index in the *i*-th data set. Each row of *P_i_* corresponds to one cell pair whose indices in the data sets constitutes the elements of *P_i_*.

Let us denote the nonlinear relationship as a function *F_i_* that maps from *R^k^* to *R^k^*, where *k* is the number of the factors in the joint NMF. The input layer of our neural network has a dimension equal to the number of factors *k*. The dimensions of seven internal layers are 200, 100, 50, 25, 50, 100, and 200 respectively. These seven layers are designed to learn and encode features that are shared across factors. Since in most data sets, the number of common factors after joint NMF is fewer than 200, we set the largest number of neurons to 200. All layers are fully connected with a rectified linear unit (ReLU) activation function, while the last layer is a fully connected layer for output ^24^. We found in practice that results were robust with respect to the chosen number of layers and numbers of neurons in each layer (Supplementary Figure 6). Throughout this study the same network architecture was utilized to analyze all data sets. The objective function of our neural network is

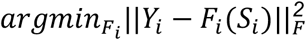

where 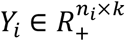 and 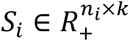 are obtained from *H_0_* and *H_i_* using the cell pair information reflected in *P_i_*. *Y_i_* is formed by the order of the index in the first column of *P_i_* from the original loading matrix *H_0_*. *S_i_* is formed by the order of the index in the second column of *P_i_* from the original loading matrix *H_i_*. The loss function is a Frobenius norm function. The objective function is optimized stochastically using Adaptive Moment Estimation with the learning rate set to 10e-5 for 500 epochs (cross-validation) ^24^.

### Co-embedding and downstream analyses

We apply UMAP or t-SNE to obtain the co-embedding plots with the inputs of combining ***H**_0_*, ***F***_1_ (***H**_1_*), ***F***_2_(***H***_2_),…, *F_N_*(*H_N_*)^25,26^. Naturally, we could use the distance ***D**_i_* (***i*** = **0,1,…,*N***) between cells from all data sets to impute the data, which is calculated as ***D_i_X_i_*** (***i*** = **1,2,…,*N***). These imputed data sets could be used to perform downstream analyses such as joint trajectory inference and finding the regulatory relationship between the cis-regulatory DNA sequences and gene promoters.

### Integration accuracy

Taking advantage of data sets that jointly profile RNA and chromatin accessibility in the same cells^7, 8, 13^, we devise a distance metric to quantify the integration accuracy of the algorithms. First, we compute the co-embedded data *C_Seurat_,C_Liger_,C_Gluer_* using each method. We use the first 50 principal components from principal component analysis in Seurat and the factor loading matrices in LIGER and GLUER. Next, we construct pairwise Euclidean distance matrices among all cells, *E_Seurat_,E_Liger_,E_Gluer_* based on coembedded data, *C_Seurat_,C_Liger_,C_Gluer_*. For each cell *a*, we calculate the integrating accuracy, *1 - ID* (*a, a*’), as the percentile of its true paired cell in its neighborhood defined using the co-embedded data, that is, *ID*(*a,a*’) = %(*a*’,*E_i_*(*a*)), where %(*a*’, *E_i_*(*a*))) is the percentile ranking of *a*’ in the neighborhood of cell *a. E_i_*(*a*) is the ranked distance vector of cell *a* to the rest of cells in the combined reference and query data sets. The greater the integrating accuracy, the better the integration result.

## DATA AND SOFTWARE AVAILABILITY

GLUER is freely available as a Python package: https://github.com/tanlabcode/GLUER. Sources of public data sets used in this paper are summarized in Supplementary Table 1.

## Acknowledgements

We thank the Research Information Services at the Children’s Hospital of Philadelphia for providing computing support. This work was supported by National Institutes of Health of United States of America grants CA233285, CA243072, and HL156090 (to K.T.), a grant from the Leona M. and Harry B Helmsley Charitable Trust (2008-04062 to K.T.) and a grant from the Alex’s Lemonade Stand Foundation (to K.T.)

## Author contributions

TP and KT conceived and designed the study. TP and KT designed and TP implemented the GLUER algorithm. TP performed data analysis. KT supervised the overall study. TP, GC, and KT wrote the paper.

## SUPPLEMENTARY MATERIALS

**Supplemental Table 1.**
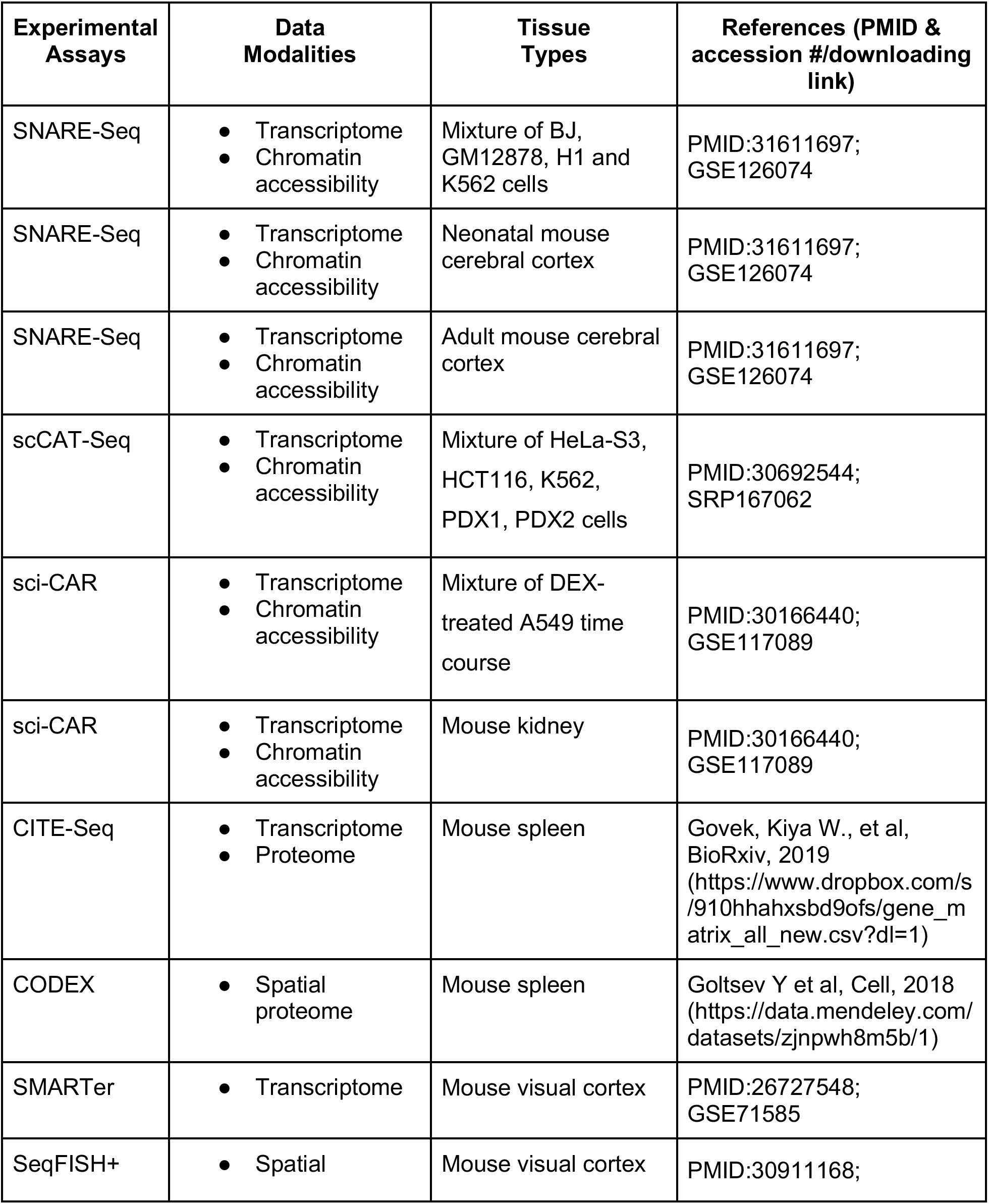
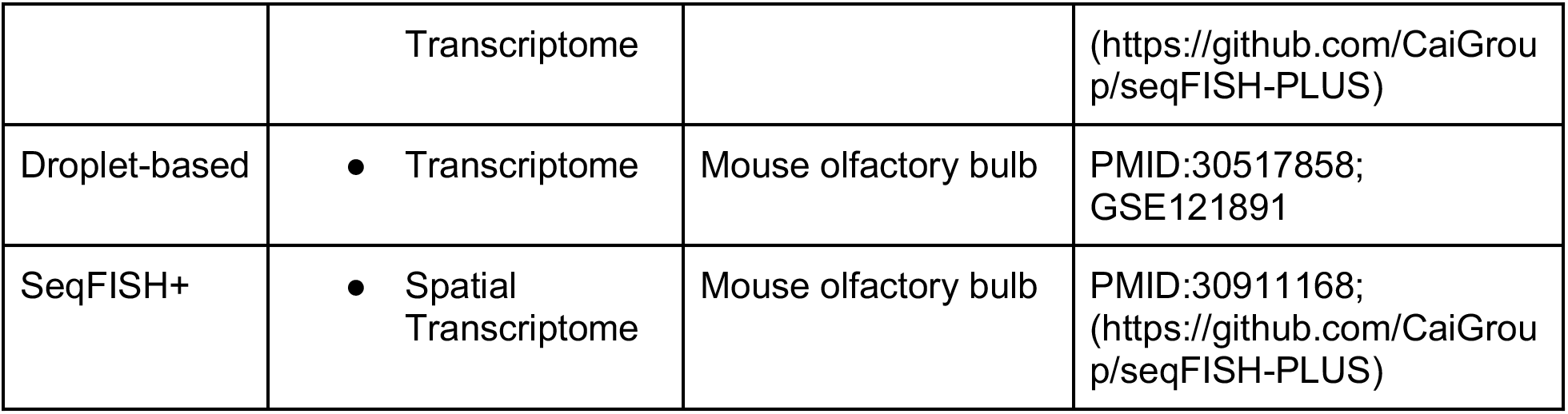
Public single-cell sequencing and imaging data sets that were used in this study.

## SUPPLEMENTARY FIGURE LEGEND

**Supplementary Figure 1.** Performance evaluation using data on a mixture of cell types profiled using the scat-Seq assay. The mixture consists of HeLa-S3 cells, HCT116 cells, K562 cells, cells derived from a moderately differentiated squamous cell carcinoma patient (PDX1), and cells derived from a large-cell lung carcinoma patient (PDX2). (**a-c**) UMAP plots of integrated data and colored-coded by data modality. (**d-f**) UMAP plots of integrated data and color-coded by cell type. (**g**) Integration accuracy of the three methods. P-values were computed using paired Student’s *t*-test with Bonferroni correction. (**h**) Distributions of gene expression levels in the original scRNA-Seq data and data computed by the integration methods. Y-axis, normalized gene expression level. *ESRP1, NPR3, SAMSN1, KRT16, LMO7* are marker genes for HCT116, HeLa-S3, K562, PDX2, PDX1 cells, respectively.

**Supplementary Figure 2.** Performance evaluation using data on DEX-treated A549 cells profiled using the sci-CAR assay. Cells were treated at three time points, 0, 1 and 3 hours before profiling. (**a-c**) UMAP plots of integrated data and color-coded by data modality. (**d-f**) UMAP plots of integrated data and color-coded by experimental condition. (**g**) Integration accuracy of the three methods. P-values were computed using paired Student’s *t*-test with Bonferroni correction.

**Supplementary Figure 3.** Performance evaluation using data on neonatal mouse cerebral cortex. 5,081 cells were profiled using the SNARE-Seq assay. (**a-c**) UMAP plots of integrated data and color-coded by data modality. (**d**) Integration accuracy of the three methods. P-values were computed using paired Student’s *t*-test with Bonferroni correction. (**e**) Cell type annotation using GLUER integration result. (**f**) Marker gene expression profiles of cell types identified in panel e. (**g**) Marker gene activity profiles of cell types identified in panel e. Y-axis, cell types, x-axis, marker genes. Shade of the dots is proportional to normalized gene expression (for RNA-Seq data) or gene activity (for ATAC-Seq data). Size of the dots represents the percentage of cells with nonzero expression or activity of the gene.

**Supplementary Figure 4**. Performance evaluation using data on 5,916 mouse kidney cells profiled using the sci-CAR assay. (**a-c**) UMAP plots of integrated data and color-coded by data modality. (**d**) Integration accuracy of the three methods. P-values were computed using paired Student’s *t*-test with Bonferroni correction. (**e**) Cell type annotation using GLUER integration result. (**f**) Marker gene expression profiles of cell types identified in panel e. (**g**) Marker gene activity profiles of cell types identified in panel e. Y-axis, cell types, x-axis, marker genes. Shade of the dots is proportional to normalized gene expression (for RNA-Seq data) or gene activity (for ATAC-Seq data). Size of the dots represents the percentage of cells with nonzero expression or activity of the gene.

**Supplementary Figure 5.** Gene/protein expression profiles of cells in mouse spleen.

(**a**) the gene expression (**b**) the protein expression. Y-axis, cell type. X-axis, gene/protein. The color is the normalized expression levels of genes computed by max-min method. The size of the dot is the percentage of cells with high expression levels of the gene.

**Supplementary Figure 6.** Performance robustness against the number of neurons in each layer of the deep neural network. X-axis, layers of the neural network; Y-axis, coefficient of variance of integration accuracy. We calculated the integration accuracy using varying numbers of neurons in each layer, including 40, 40, 20, 10, 20, 40, and 40 different numbers of neurons for layers 1 to 7, respectively.

